# A Growing Microcolony can Survive and Support Persistent Propagation of Virulent Phages

**DOI:** 10.1101/149062

**Authors:** Rasmus Skytte Eriksen, Sine Lo Svenningsen, Kim Sneppen, Namiko Mitarai

## Abstract

Bacteria form colonies and secrete extracellular polymeric substances that surround the individual cells. These spatial structures are often associated with collaboration and quorum sensing between the bacteria. Here we investigate the mutual protection provided by spherical growth of a monoclonal colony during exposure to phages that proliferate on its surface. As a proof of concept we exposed growing colonies of *Escherichia coli* to a virulent mutant of phage P1. When the colony consists of less than ~ 50000 members it is eliminated, while larger initial colonies allow long-term survival because the growth of bacteria throughout the spherical colony exceeds the killing of bacteria on the surface. A mathematical model pinpoints how this critical colony size depends on key parameters in the phage infection cycle. Surprisingly, we predict that a higher phage adsorption rate would allow substantially smaller colonies to survive a virulent phage.

**Significance Statement:** Bacteria are repeatedly exposed to an excess of phages, and carry evidence of this in terms of multiple defense mechanisms encoded in their genome. In addition to molecular mechanisms, bacteria may exploit the defense of spatial refuges. Here we demonstrate how bacteria can limit the impact of a virulent phage attack by growing as a colony which only exposes its surface to phages. We identify a critical size of the initial colony, below which the phages entirely eliminates the colony, and above which the colony continues to grow despite the presence of phages. Our study suggests that coexistence of phages and bacteria is strongly influenced by the spatial composition of microcolonies of susceptible bacteria.

---

Virulent phages are ubiquitous, yet seemingly at odds with their own survival. As they kill their susceptible hosts, they suppress these to low populations levels (1–3), creating a fragile and hugely competitive environment for themselves (e.g. (4–6)). By contrast, temperate phages have the option to lysogenize their host and thereby preserve their DNA through periods of low host availability (7). This apparent advantage of temperate over virulent phages suggests the need for a more fine-grained understanding of virulence.

Noticeably, the virulent phage species present a quite diverse set of characteristics for their killing. Even under identical conditions, different species have average burst sizes that vary by more than a factor 10 and have latency times that vary by a factor four (8). Phage T4 extends this latency period when there are several infecting phages in a host simultaneously, a phenomenon known as lysis inhibition (9, 10), and also has the ability to enter a prolonged “hibernation” mode after initiation of progeny phage protein synthesis in starving cells (11). Some virulent phages such as T3 and T7 are known to form particularly large plaques (12), while for example T4 forms rather small plaques due to lysis inhibition (9, 12). The variation in the “art of killing” is also indicated by the fact that some virulent phages (e.g. *Qβ*) encode less that ten genes(13), while others (e.g. T4) encode more than double the gene number of the temperate phage λ (14,15).

Here we explore a new aspect of virulence, namely how a virulent phage may propagate in a growing bacterial colony. Our investigation is inspired by visual inspection of the plaque formed by a virulent phage in Fig. 1. The picture shows a plaque that is typically categorized as a “clear plaque”, started by one P1_Vir_ phage that was introduced onto a bacterial lawn consisting of *Escherichia coli* cells. A nearly clear region surrounds the initial infection at the centre, but one can also observe faint colonies that are larger near the edge of the plaque. Outside the plaque, the agar contains a dense collection of colonies that have grown to stationary phase without the influence of phages. The center of the plaque is where the first infection occurred, whereas bacteria in the periphery of the plaque encountered phages relatively later, after each bacterial cell had grown to form a microcolony (16). The origin of the faint colonies at the plaque periphery has not been fully understood (17).

**Fig. 1.**
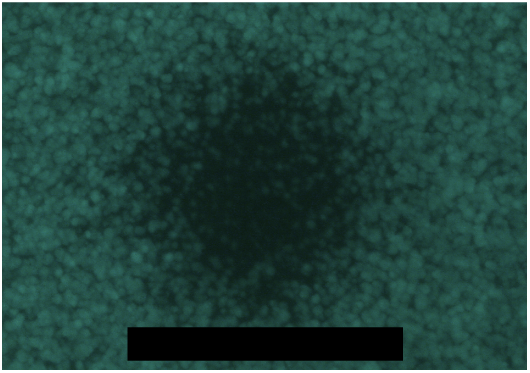
Plaque formed by P1_vir_. Green spheres are microcolonies of *E. coli* expressing green fluorescent protein. Notice the faint colonies, which are seen across the plaque although less in its central region. Scale bar: 300 *μ*m.

This paper explores the fate of growing bacterial colonies that are exposed to phages at various time-points during their growth. A previously proposed scenario (17) discussed that cells in the center of a large microcolony may enter stationary phase and become unable to support the production of progeny phages. Though such a protection mechanism is certainly expected, a recent work showed that the growth of even quite large *E. coli* colonies ( ~ 10^7^ cells) mimics the normal exponential growth of individual *E. coli* cells in liquid medium (18). This finding indicates that nutrients diffuse quite extensively through the colony. By contrast, phages may well adsorb to cells on the colony surface and therefore only rarely reach deeper layers of the colony. Thus, we hypothesize that the continued growth from the inside of a sufficiently large microcolony could potentially overwhelm phage-mediated killing on the microcolony surface. In combination with the phage-refractory state of stationary phase cells, the result would be long-term survival of phage-sensitive cells inside the microcolony.

Here, we first construct a model of growing cells and infecting phages to numerically simulate the phage attack on a growing microcolony. The simulation suggests a sharp transition from elimination of the microcolony to its continued growth as the microcolony size at the time of the first phage encounter becomes larger. We then show that the model prediction is consistent with an experiment by infecting *E. coli* microcolonies of different sizes with P1_vir_ and monitoring their fate. Our study demonstrates that a sufficiently large microcolony indeed provides a refuge against virulent phages. The refuge counteracts the phages’ ability to control bacterial biomass (6), and thereby allows the bacteria to circumvent the famous kill-the-winner (19) feature of phage predation in well-mixed environments.

## Results

### Modeling a phage attack on a growing microcolony

We simulate microcolony growth by modeling each cell as a sphere, and each cell grows and divides exponentially with a maximum doubling time of 20 minutes. The repulsive force between the cells provides an overall spherical growth of the microcolony consisting of densely packed bacteria. At a certain time, we let the surface of a grown microcolony be surrounded by ~ 2700 phages, and observe whether the phages kill all the cells or if the cells can outgrow the ongoing lysis by the phages. The simulated microcolonies are rather small, and the explicit modeling of nutrient gradients inside a microcolony gave only minor growth rate differences across the colony (Supporting Information). Therefore we here present the results from simulations that assume a constant nutrient level across the microcolony. The detailed simulation protocol is described in the Methods section and Supporting Information, and the default parameter values are listed in Supporting Table S1.

Fig. 2 illustrates model simulations with healthy bacterial cells shown as blue spheres. Bacteria infected by phages are colored from yellow to red, where the ones closest to lysis are assigned the most reddish color. Each lysis event is modeled by replacement of the bacterium with 400 point-like phage particles at the end of the latency period. The progeny phages (not shown) are allowed to diffuse and infect other bacteria. In this simulation, we assume that a phage particle adsorbs to a cell as soon as they overlap, mimicking the diffusion-limited scenario of adsorption. The phages adsorb equally well to uninfected cells and those that are already infected by another phage(s). We later study the effect of a smaller adsorption rate on microcolony growth.

**Fig. 2.**
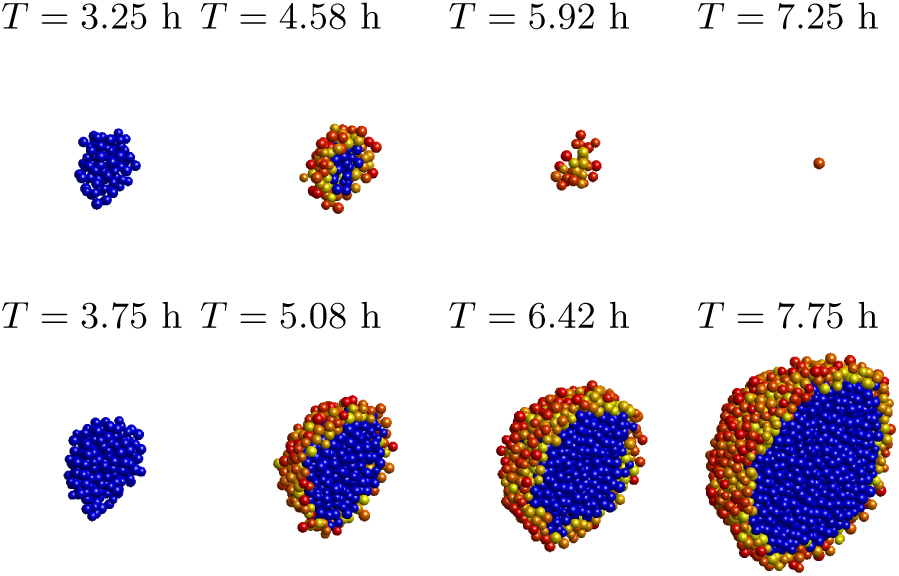
Snapshots of simulations with half of the microcolonies presented. The upper panel shows the case of a relatively early phage attack (infection at 3.25h), which results in elimination of the microcolony. The lower panel follow the history of a slightly later attack (infection at 3.75h), which results in continued growth of the microcolony.

In the upper row of panels in Fig. 2 the first attack occurs at 3.25 hours of growth, when the colony has a size of 220 members. The lower panels depict a phage attack at 3.75 hours where the colony size has reached 500 members. One sees that the smaller colony is eventually entirely eliminated, which in turn would leave the fate of the phage particles to the chance that they reach another colony before they decay. In contrast, when the phages attack the larger colony it survives and grows with phages persisting on the infected surface. The simulation showed that the distance phages can penetrate into the microcolony, ΔR, is roughly constant when the microcolony size is above the threshold of killing. For the parameters used in Fig. 2, phages can infect about 2 layers of cells. This is because, when the infected cells at the surface lyse, the phage progeny mostly adsorb to the cells around them before they can diffuse deeper into the microcolony.

#### A minimal critical radius for survival

Fig. 3 examines the predictions of the model. Panel A) shows the final growth rate of a colony as a function of the time at which it first encountered the phages. One sees that our model indeed predicts a sharp transition from extinction to persistent growth, when the size of the microcolony have surpassed a lower critical threshold. One also sees that when the first infection is sufficiently late to allow colony survival, then the colony will grow to a size where the growth penalty associated with infections at its surface becomes relatively small. Notice that the growth rate is measured while the nutrient depletion is assumed to be negligible. In reality this assumption will ultimately break down when the inner part of the colony becomes increasingly shielded from available nutrients. The lack of nutrients will however also prevent or at least reduce phage proliferation. Thus, if the microcolony is above the survival threshold, it is predicted to eventually maintain a large number of living sensitive cells in the center.

**Fig. 3.**
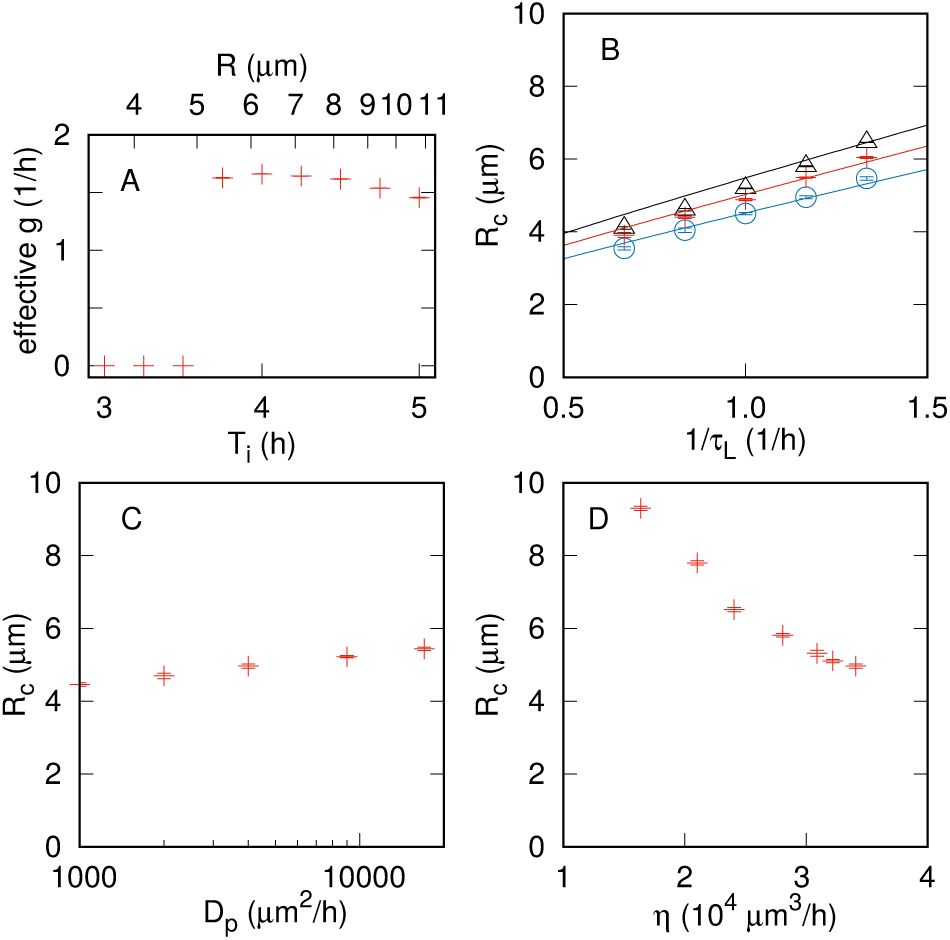
Model predictions. **A.** Long-term growth rate of phage-infected microcolony as function of initial exposure time (*T_i_*). The effective growth rate *g* was determined by fitting to the analytical solution eq. (S8) of the linearized version of the model equation to the simulation time traces of the radius *R*(*t*) (see Supporting Information for detail). The radius of the colony at the initial exposure time can be read on the top horizontal axis. **B.** The critical radius *R_c_* as function of latency time *τ_L_* for burst sizes of 100 (open blue circles), 400 (red crosses), and 1000 (open black triangles). The fit with eq. (S7) to each data set is shown as a solid line, with the fitting parameter Δ*R* =1.26,1.40,and 1.53 *μ*m, respectively. **C.** *R_c_* as a function of the phage diffusion constant *D_P_*. **D.** *R_c_* as function of the phage adsorption rate *η*. The standard error of the mean is shown as error bars.

Assuming a constant penetration depth Δ*R* of the phages into a growing colony allows us to understand the steep size-dependent transition in microcolony fate. Suppose that a spherical microcolony of radius *R*(*t*) at time *t* consists of an inner core of exponentially growing uninfected cells with growth rate of biomass *g* and an infected surface layer of a constant thickness Δ*R*. We further assume that infected cells will burst with a rate 1/*τ_L_*, with *τ_L_* being the latency time of the phage burst. This leads to the equation for the total volume 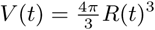 to be

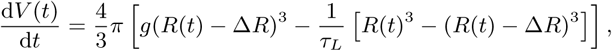

where the first term in the brackets depicts the growth of the uninfected cells in the inner core, while the second term describes the lysis of the outer infected layer. Clearly, the growth of the total volume d*V* (*t*)/d*t* > 0 requires large enough *R*(*t*) compared to Δ*R*. The critical threshold *R_c_* for the initial size *R*(0) is determined by d*V* (0)/d*t* = 0, and the solution is given in the supplement eq. (S7) that fits well with the simulation (Fig. 3B). If we approximate the solution for a small penetration depth case (Δ*R* ≪ *R_c_*), we get *R_c_* = 3[1 + (*gτ_L_*)^−1^]Δ*R*, depicting that *R_c_* increases with Δ*R*, and that a smaller burst rate 1/*τ_L_* compared to the cell growth rate *g* decreases *R_c_*.

The critical radius *R_c_* depends on the various parameters as demonstrated in Fig. 3B-D. *R_c_* grows roughly linearly with the burst rate 1/*τ_L_* (Fig. 3B), because the latency time sets the timescale for the lysis of cells next to the uninfected core. The burst size affects Δ*R* and hence *R_c_*, but changing the burst size from 100 to 1000 only increases the critical radius *R_c_* by about one cell layer with the given parameter set (Fig. 3B). A larger diffusion constant *D_p_* only marginally increases the critical radius (Fig. 3C), because the phages are hindered from reaching the core cells by adsorption to the infected surface cells regardless of the magnitude of the diffusion constant.

Reducing the adsorption rate *η* from the diffusion-limited case causes a substantial increase in the critical radius *R_c_*. To simulate small *η* we allowed the phage particles that overlap a cell to be repelled away and introduced a finite rate *γ* with which infection can occur while there is overlap. Reducing *γ* parameterizes a reduction of the phage adsorption rate *η* from the diffusion-limited value. The reduction of adsorption rate *η* dramatically increases the critical radius *R_c_* (Fig. 3C). This reduction is caused by a increased penetration depth Δ*R* as phages can diffuse deeper into the colony core before they adsorb to a cell.

### Infection of *E. coli* microcolonies with P1_vir_

To test the above predictions in the laboratory, we performed infections of *Escherichia coli* microcolonies with a virulent mutant of the phage P1, P1_vir_ (20). This phage has a fairly long latency time of 60 min for *E. coli* growing in rich medium (8), thus facilitating a moderately small value of *R_c_*. As the bacterial host, we used *E. coli* strain SP427 (21), which expresses green flourescent protein (GFP) constitutively. The GFP expression allowed us to visualize the intact cells in the colonies using fluorescence microscopy. To obtain approximately spherical microcolonies, we grew the host cells embedded in a thin soft-agar layer on plates containing rich medium, and sprayed these plates with phage lysate at different times after incubation, thereby implementing phage attacks on microcolonies of different sizes. The detailed experimental protocol is given in Methods.

Fig. 4 shows images of representative colonies at the time of addition of phage P1_vir_, and at 16 hours of incubation post phage addition. The left panels show the dark-field images, whereas the right panels show the green fluorescence images, which visualize intact bacteria that contain GFP. The panel “5h” displays the rather small colony size that was typically obtained after 5 hours of incubation. The “5h+16h” panel at the top right shows the typical final colony after an additional 16 hours of incubation without exposure to phage. In this case, the colony is not only large but also dense and homogeneously fluorescent. In contrast, the middle-right panel “5h+16h phage” shows a typical final colony when the plate has been sprayed with phage lysate after 5 hours of growth and then incubated 16 hours after that. One sees that the colony has grown bigger after incubation in the presence of phages, but it exhibits only faint fluorescence, indicating that most of the colony shown in the dark-field image consisted of dead remnants of bacteria.

**Fig. 4.**
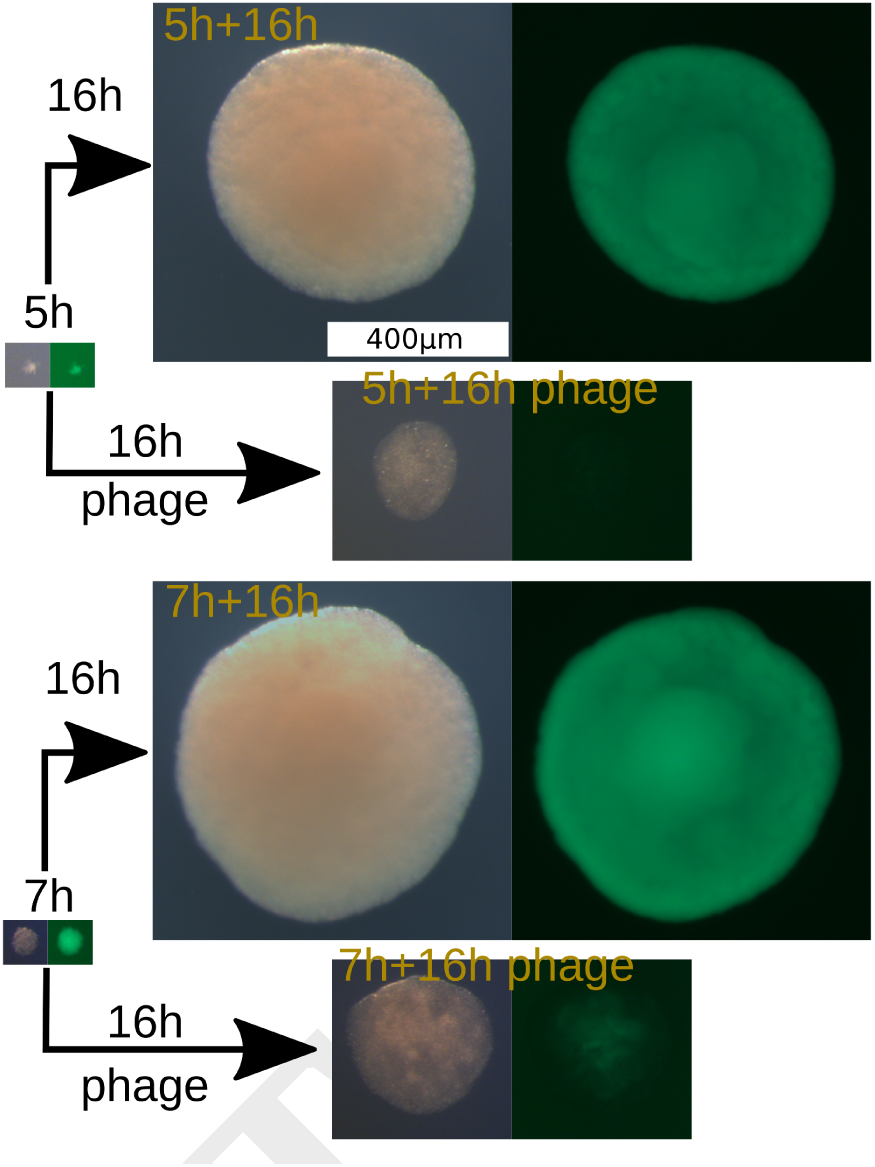
Images of typical microcolonies before and after exposure to phages. The dark-field image and the corresponding green fluorescence image of the same colony are shown side by side. The green fluorescence images in the final colonies with phage exposure at 5h and at 7h spray time were collected at the same light source strength and with the same exposure time to allow direct comparison.

The typical colony obtained after 7 hours undisturbed growth is shown in the bottom “7h” panel, and is visibly larger than the typical colony after 5 hours of incubation, reflecting the additional four doublings of biomass. The panel “7h+16h phage” shows that the final size of this colony after they were exposed to phages and then incubated for 16 more hours. Though it is visibly smaller than the undisturbed colony in “7h+16h” panel, one observes a fluorescent region in the middle, suggesting a substantial amount of surviving bacteria.

The colony growth is deduced from microscopy images. Fig. 5A shows that the radius grows exponentially when there is no exposure to phages. This growth is consistent with a doubling time of ~ 30 minutes for about 9 hours until a size of ~ 2 × 10^6^ bacteria if we assume a spherical colony of dense cells with volumes of 1 *μ*m^3^. After about 10 hours, the growth rate is reduced substantially and colony size eventually stabilizes at a level set by the initially available nutrients and the number of colonies per plate sharing those nutrients. Fig. 5C shows how this final colony size varies moderately between experiments without phage exposure. The final radius of 400*μ*m implies that our conditions allow colonies to reach ~ 2·10^8^ bacteria. Because slow-growing bacteria are smaller than rapidly growing bacteria (22), we cannot accurately deduce the actual number of bacteria in these colonies.

**Fig. 5.**
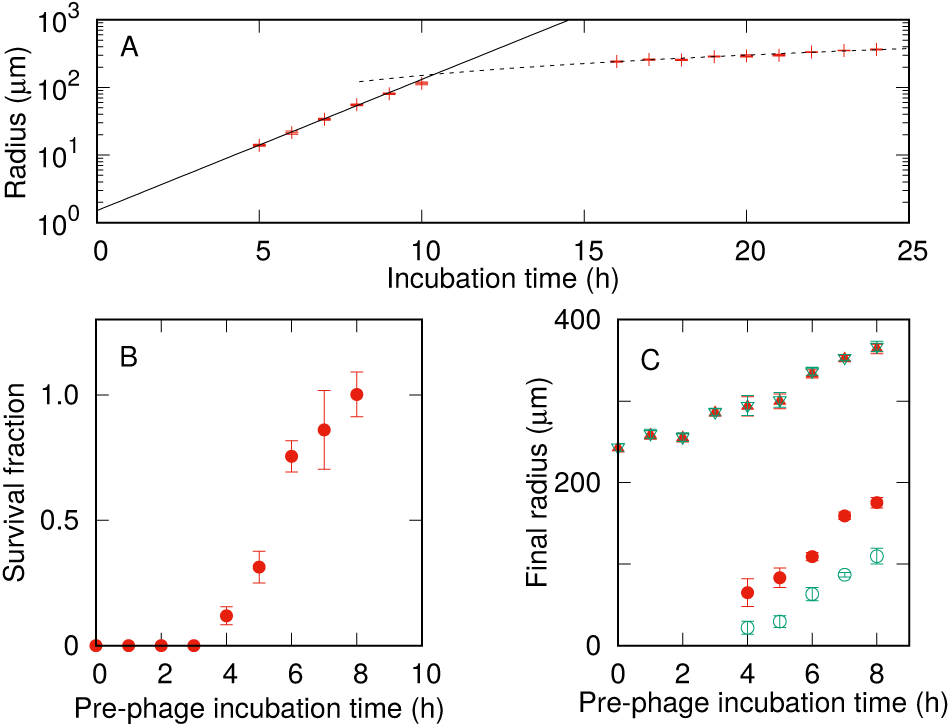
P1_vir_ infection of *E. coli* microcolonies. **A**. Growth in colony radius over time in the absence of phages. The solid line marks an exponential fit with a doubling time of 31 min. The dashed line is a linear fit to growth in colony radius on longer timescales. The cross-over point is ~ 10 hours. The radius was estimated from the colony image from the area *A* as 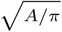. **B**. Number of visible colonies in the plates sprayed with phages relative to the mean number of colonies on control plates without phages. The control plates had 76±4 visible colonies per plate. The horizontal axis shows the microcolony growth time prior to the exposure to phages. **C**. Final colony size as a function of the duration of the pre-phage incubation. The filled red symbols show the estimates from dark-field images whereas the open green symbols show the radius estimated from the green fluorescence images. The triangle symbols between 200-400 *μ*m show the final radius of colonies in the absence of phages, while the round symbols show the final radius of colonies exposed to phages. The standard error of the mean is shown as error bars.

Predictably, exposure to phages alters the colonies dramatically. Fig. 5B quantifies the number of colonies per plate that are visible with the naked eye after the final incubation period for different pre-phage incubation times. The surviving fraction is normalized to the average number of visible colonies per control plate that was not exposed to phage. Almost no colonies had grown to visible size when the phages were introduced earlier than 4 hours of incubation. The absence of surviving colonies was confirmed using fluorescence microscopy with 10X magnification. After this critical time, visible colonies emerged. When phages were introduced at 4 hours, a few small colonies could be detected. When phages were introduced at 6 hours and later, almost 100% of the initial cells grew to form visible colonies. Thus, this laboratory experiment reproduced a steep transition in the number of surviving colonies that depended on the size of the microcolony at the time of phage addition. Two additional repetitions of the experiment confirmed the jump in colony survival when phages were introduced after 5-6 hours of pre-phage incubation (see Supporting Information for details).

Fig. 5C shows the final size of colonies that were exposed to phages (round symbols). In the dark-field recording we see a moderate change in the radius of final colonies from about 90 *μ*m to about 180 *μ*m as the phage exposure time is delayed. The corresponding change in size of the GFP expressing part of the colony increases more dramatically from 20 *μ*m to about 100 *μ*m. In all cases, the outer layers of the colony appear dark, indicating that the surfaces of these colonies consist of bacterial debris from *E. coli* that were killed by phages.

An alternative explanation for the microcolony-size-dependent jump in colony survival (Fig. 5B) could be that the surviving cells are all resistant to P1_vir_ infection, and the jump reflects the critical colony size at which there is a large probability that the colony contains one or more resistant mutants at the time of phage exposure. To distinguish between this model and the spatial refuge model, we first tested if the colonies contain surviving cells, and if there were any, we tested whether they are sensitive or resistant to P1_vir_. The overnight culture used for the experiment depicted in Fig. 5 contained P1_vir_ resistant mutants at a frequency of 1.2 × 10^−5^ consistent with a previous measurement (23).

We picked 10 colonies from each of the phage-sprayed plates that had experienced at least 4 hours pre-phage incubation time, and streaked them on fresh plates containing the chelator sodium citrate to inactivate free phages and thereby permit colony formation by both sensitive and resistant survivors from the sampled colonies. As summarized in Table 1 (# recovered), only some of the sampled colonies gave rise to new colonies on the fresh plates, confirming survival after long-term phage exposure. We note that the lack of colony formation does not necessarily mean that all cells in a sampled colony were dead, because we picked cells from the soft-agar-embedded colonies with a toothpick, rather than extracting and plating whole colonies. From each re-streak, 10 individual colonies (or all the colonies if there were less than 10) from each plate were tested for P1_vir_ resistance by cross-streaking (resulting in cross-streaking of colonies corresponding to 100 cells from each pre-phage incubation time point, provided there was sufficient growth on the re-streaks). In the results summarized in Table 1 (# with resistance), an original colony was counted as resistant if any of the up to 10 resulting colonies were tested positive for resistance. The detailed results of the resistance test are shown in Table S2. These results show that although there were naturally a substantial number of resistant cells in the surviving colonies, we did not find any resistant cells in about half of the sampled colonies, indicating that the majority of the survivors in those colonies were sensitive to P1_vir_.

**Table 1.** P1_vir_ resistance test summary. Details are in Table S2.

| Time | # restreaked | # recovered | # with resistance | # sensitive |
| --- | --- | --- | --- | --- |
| 4h | 10 | 6 | 2 | 4 |
| 5h | 10 | 5 | 1 | 4 |
| 6h | 10 | 7 | 5 | 2 |
| 7h | 10 | 9 | 5 | 4 |
| 8h | 10 | 10 | 5 | 5 |

## Discussion

This paper argues for a new type of herd-immunity associated with consecutive layers of bacteria that shield each other from infection. We modeled colonies exposed to phages at various time-points in their growth, and pinpointed that the latency time and the adsorption rate should be particularly important parameters for deciding the fate of the infected colony. Smaller adsorption rates increase killing of the host colony, since phages penetrate deeper into the microcolony without being adsorbed by the infected cells on the surface. A similar effect was observed previously for phages infecting bacteria in biofilm (24), indicating the importance of optimization of adsorption rate for phages to spread in a spatially structured habitat.

We experimentally demonstrated that shielding can take place, by showing that sensitive cells can constitute the majority of the survivors in a growing phage-infected colony. Experimentally, the critical microcolony growth time is not as sharp as in the numerical simulation. This is expected because the vertical position of the microcolonies vary within the ~ 0.4mm thick top-agar layer, and the colonies at the bottom will be exposed to phages later than those nearer the top.

Indeed, we confirmed in additional experiments that the soft-agar thickness is an important factor, but the jump of the survival fraction to almost 100% at around 6 hours pre-phage incubation time was confirmed, also in experiments where we varied the soft agar thickness (See Supporting Information for details). The additional time needed for diffusion through the top-agar brings a conservative estimate of the actual critical phage arrival time to about 6 to 7 hours, resulting in an estimated *R_c_* ≈ 25*μ*m critical microcolony radius. Image analysis confirmed that each final colony had grown significantly beyond this size, supporting the proposed survival scenario. Quantitatively, this *R_c_* value is about 3 times larger than the typical results of the simulation in Fig. 3. Part of this disagreement could be because we simplified the cell shape to be a sphere in the simulations, while *E. coli* is known to have an elongated rod shape. If the long-edge of a cell is pointed vertical into the microcolony, as is apparently the norm for other rod-shaped bacteria (25) the burst of such a cell allow phages to penetrate significantly deeper than in our scenario. Thereby the elongated shape of *E. coli* increases the penetration depth Δ*R*, and hence increases the critical radius *R_c_* proportionally.

The fate of a colony may not only depend on whether its growth can outrival the phage-mediated deaths on the surface. In particular, when bacteria grow slower, phage infections tend to yield smaller burst sizes with longer latency time (26–28). This means that a colony which approaches its resource-limited size may survive simply because the phages cannot propagate. In our study, this effect was not important for the determination of the critical threshold, since the threshold for colony survival was at most 7 hours, and exponential growth continued past the 9 hour mark. Thus, the surviving colonies had certainly grown after encountering the phages. Naturally, for survival of sensitive cells after overnight incubation with phages, it is also important that the phages could not propagate on stationary-phase cells.

Theoretical models supplemented with experiments on phage-bacteria culture in liquid media demonstrated (1, 2, 19) that bacteria and virulent phages persistently can co-exist. They do so in self-organized critical conditions where only one phage from each bacterial lysis event persists to infect another susceptible host. Our present study suggests to view this coexistence differently when dealing with bacteria growing in a gel or a semi-solid, as for example in soil. Because all new potential hosts are generated from bacteria that are already present, the spatial distribution of the host will be maintained over long time periods. The destiny of released phages then become crucially dependent on the distances to new colonies of hosts, and the overall survival of a virulent phage may then rely on “farming” the local colony in a sustainable way.

The observation of bacterial colonies with an active core surrounded by an extended region of likely dead (non-fluorescent) bacteria suggest yet another possibility that may enhance colony robustness. As dead bacteria accumulate on the surface of the colony, phage particles may lose their DNA by injecting it into these hosts remnants. Thus, the debris may cause a sink for phage DNA at the surface of the colony that further enhances survival in its core.

In any case, our paper suggests that the evolution of especially long latency time and high adsorption rate for a virulent phage may reflect its ability to live in a persistently infectious parasitic state on a growing bacterial colony. A related persistent infection pattern is seen in filamentous phages (e.g. M13) that, on the scale of the individual bacteria, maintains the host alive while producing phages (29). In contrast, phages with small adsorption rates and short latency times will more easily exterminate a colony. One outcome of our colony-focused “micro-cosmos” is thus that the “choice” of latency time for a phage is much more than just an optimization between fast generations with few off-springs and longer generations with much larger burst size of phages (30). For lytic phages, the “art of killing” (31) has more facets than growth on a well-mixed host population.

## Methods

### Outline of the numerical simulation

Cells are modeled as a 3-dimensional spheres that follow the over-damped dynamics and repel each other when they overlap. An uninfected cell grows exponentially with a nutrient-dependent rate, and divides into two cells when it reaches a threshold volume. We set the parameters so that the median of the cell volume is 1.3 *μm*^3^ and the maximum cell generation time is 30 min. We assume nutrients are spatially homogeneous and consumed as the cells grow. A phage is modeled as a point particle that obeys the over-damped Langevin equation. When the phage overlaps with a cell, a repulsive force is exerted from the cell. A phage overlapping with a cell is adsorbed with a rate *γ*, where *γ* = ∞ models the diffusion limited adsorption, while the phage adsorption rate *η* can reduced by limiting the *γ* value. When a cell is infected by a phage, it produces phage particles after an average latency time *τ_L_*, modeled as a sequence of 10 Poisson processes (16). During the latency period, an infected cell can adsorb more phages. When a lysis occurs, *β* new phage particles are spawned uniformly around the cell’s center and the cell is removed. The simulation starts with a single cell, which is allowed to establish a colony for the time duration *T_i_*. At time *T_i_* phages are spawned uniformly in the simulated volume outside of the colony. More details are given in Supporting Information.

### Experimental procedures

#### Infection of E. coli microcolonies with P1_vir_

*E. coli* strain SP427 (21), a MC4100-derivative containing a chromosomally encoded *P_A_*_1/*O*4/*O*3_::*gfp*mut3b gene cassette (32), was grown overnight in LB (33) supplemented with 50 *μ*g/ml kanamycin. The culture was diluted to ~ 400 CFU/ml in sterile MC buffer (50 mM CaCl2, 25 mM MgCl2) and kept at room temperature. Every hour, 250 *μ*l of the diluted culture was mixed with 2.5 ml R-top agar (as (34) but with 5 g/L agar and 50 *μ*g/ml kanamycin), plated on fresh R agar plates (as R-top but with 12 g/L agar), and incubated at 37°C to allow individual cells to form microcolonies. After a given incubation time at 37°C, 0.5 ml of P1_vir_ phage lysate (7 × 10^9^ PFU/ml) was sprayed uniformly on the top agar surface using a perfume bottle with atomizer bulb, and the plate was incubated for another 16 hours at 37°C. One control plate without the addition of P1_vir_ were processed along with triplicate phage-spray plates for each time point. The experiment was repeated three times with similar results. Details of the repeat experiments can be found in Supporting Information, section 4. Microscopy images were obtained using a Leica MZ16 F fluorescence stereomicroscope and quantification was done by matlab code developed by the authors. The periphral 1cm of each plate were excluded in the analysis to avoid any effects of heterogeneity in phage spraying close to the edge. Six additional plates without phage addition were prepared as the control plates above and used for microscopy imaging to determine microcolony size from 5 to 10 hours of incubation at 37°C.

#### P1_vir_ Resistance test

To determine the level of P1_vir_ resistance among cells in the culture used for seeding the microcolonies, dilutions of the culture in MC buffer were mixed with P1_vir_ at an average phage input >10, incubated for 20 min, and plated on LB plates. To test individual cells in the surviving microcolonies for whether they were sensitive or resistant to P1_vir_, 10 individual microcolonies were picked out of the top agar from a plate for each time point (4-8 hours) and restreaked on tryptone plates (34) containing 10 mM sodium citrate to destabilize the phage particles and thereby permit colony formation by both sensitive and resistant cells. The number of plates that contained *E. coli* colonies after overnight incubation is shown in Table 1 (# recovered). From these survivors, up to 10 individual colonies were picked from each plate, and cross-streaked against P1_vir_ on R plates supplemented as above. The outcome of the cross-streak was confirmed for 36 colonies by infection with P1_r*ev*6_ (35) and selection for lysogeny on LB plates containing 15 *μ*g/ml chloramphenicol.

## ACKNOWLEDGMENTS

Authors sincerely thank Stanley Brown for helpful discussions. This work was funded by Danish National Research Foundation (BASP: DNRF120).

